# The kinase specificity of protein kinase inhibitor peptide (PKI)

**DOI:** 10.1101/2020.11.25.399204

**Authors:** Yao Chen, Bernardo L. Sabatini

**Affiliations:** Department of Neuroscience, Washington University School of Medicine, St. Louis, MO 63110, USA; Howard Hughes Medical Institute; Department of Neurobiology, Harvard Medical School, Boston, MA 02115, USA

**Keywords:** protein kinase A, Protein kinase C, Protein kinase inhibitor peptide, kinase screen, specificity, endogenous, inhibition, facilitation

## Abstract

G-protein-coupled-receptor (GPCR) signaling is exquisitely controlled to achieve spatial and temporal specificity. The endogenous protein kinase inhibitor peptide (PKI) confines the spatial and temporal spread of the activity of protein kinase A (PKA), which integrates inputs from three major types of GPCRs. Despite its wide usage as a pharmaceutical inhibitor of PKA, it was unclear whether PKI only inhibits PKA activity. Here, the effects of PKI on 55 mouse kinases were tested in *in vitro* assays. We found that in addition to inhibiting PKA activity, both PKI (6-22) amide and full-length PKIα facilitated the activation of multiple isoforms of protein kinase C (PKC), albeit at much higher concentrations than necessary to inhibit PKA. Thus, our results call for appropriate interpretation of experimental results using PKI as a pharmaceutical agent. Furthermore, our study lays the foundation to explore the potential functions of PKI in regulating PKC activity and in coordinating PKC and PKA activities.

## Introduction

G-protein-coupled receptor (GPCR) signaling is highly regulated, and its exquisite spatial and temporal specificity has important implications on its function. Spatially, GPCRs and their signaling components are located in specific cell types, subcellular compartments, and microdomains, and their spatial localization specifies their functions (Smith et al., 2006; Chen and Sabatini, 2012; Lur and Higley, 2015; Jong et al., 2018; Thomsen et al., 2018; Lobingier and von Zastrow, 2019; Weinberg et al., 2019). Temporally, GPCR activation can lead to transient, sustained, or oscillatory patterns of intracellular signals, and the timing of GPCR activation relative to synaptic inputs is critical for how synapses are modified in the nervous system (Gu and Yakel, 2011; Muñoz and Rudy, 2014; Yagishita et al., 2014; Grundmann and Kostenis, 2017). Thus, understanding the regulators of the spatial and temporal features of GPCR signaling is important to understand GPCR functions.

One important intracellular integrator that lies downstream of multiple GPCRs is protein kinase A (PKA) (Gilman, 1995; Chen et al., 2017). PKA is activated by cyclic AMP (cAMP), which is produced by adenylate cyclases (ACs) (Krebs et al., 1959; Sutherland et al., 1968; Gilman, 1995). AC activity is stimulated by Gαs-coupled receptors and inhibited by Gαi-coupled receptors (Gilman, 1995). Furthermore, we recently discovered that endogenous Gαq-coupled receptors also activate hippocampal PKA (Chen et al., 2017). Therefore, three out of the four classes of GPCRs converge to regulate PKA activity. Furthermore, PKA phosphorylates diverse substrates to regulate synaptic and cellular functions, both within and outside the nervous system (Brandon et al., 1997; Greengard, 2001). Therefore, PKA is a biochemical integrator of signaling from numerous GPCRs that exert important cellular and physiological functions.

The spatial and temporal specificity of PKA is under exquisite control, notably by the A-kinase-anchoring proteins (AKAPs), the regulatory subunits of PKA, and the endogenously expressed PKA inhibitory peptide, PKI (Walsh et al., 1971; Taylor et al., 1990; Dalton and Dewey, 2006; Smith et al., 2006). PKI binds to the catalytic subunit of PKA, thereby preventing it from being active (Ashby and Walsh, 1972, 1973) despite release from a PKA regulatory subunit. Synaptic stimulation decreases the expression of one of the isoforms of PKI, PKIα (De Lecea et al., 1998). Furthermore, chronic infusion of antisense *pkiα* reduces neuronal excitability and eliminates hippocampal plasticity (De Lecea et al., 1998). Therefore, PKI inhibits PKA activity and has important cellular functions.

In addition to the important functions of endogenous PKI, shorter peptides of PKI, for example PKI (6-22) amide, are widely used as pharmaceutical agents to inhibit PKA activity (Cheng et al., 1986; Glass et al., 1989a, 1989b). However, it was unclear if PKI inhibits only PKA or if it inhibits other enzymes as well. Previous studies examined the effect of PKI on a few kinases, including protein kinase G (PKG) and protein kinase C (PKC) (Gill et al., 1976; Cheng et al., 1986; Glass et al., 1986; Day et al., 1989; Smith et al., 1990), but the effects of PKI on a broad panel of kinases were not known.

Here, we analyzed the effect of PKI on 55 kinases, selected based on their similarity to PKA and their expression patterns. We confirmed that both PKI (6-22) amide and full length PKIα inhibited PKA activity with sub-nanomolar IC50. In addition, we found that at high concentrations, PKI (6-22) amide inhibited calcium/calmodulin-dependent protein kinase I (CamK1). Surprisingly, at concentrations often used in pharmacology experiments, PKI (6-22) amide facilitated the activity of multiple PKC isoforms, rho-associated, coiled-coil-containing protein kinase 1 (ROCK1), and p70S6 Kinase (p70S6K). Synthesized full length PKIα also facilitated the activity of ROCK1 and multiple PKC isoforms. These results are important not only for interpretation of experiments using PKI as a pharmacological agent, but also sheds light on potential biological functions of endogenous PKI.

## Material and Methods

### PKI

PKA inhibitor (PKI) fragment (6-22) amide was purchased from Tocris (Cat. No. 1904). Full length mouse PKIα was synthesized from L amino acids and then purified with High-performance liquid chromatography (HPLC) by the Koch Institute Biopolymers and Proteomics Facility.

### Kinase Assays

Kinase assays were conducted with the Thermo Fisher Scientific SelectScreen Kinase Profiling Service. CamK1, DAPK1, and NUAK1 (ARK5) were screened with the Adapta Universal Kinase Assay, and the rest of the kinases were screened with the Z’-LYTE Peptide Kinase Assay (Rodems et al., 2002) (Figure 1B). The % inhibition value was calculated with the formula specified in the assay manuals.

**Figure 1.**
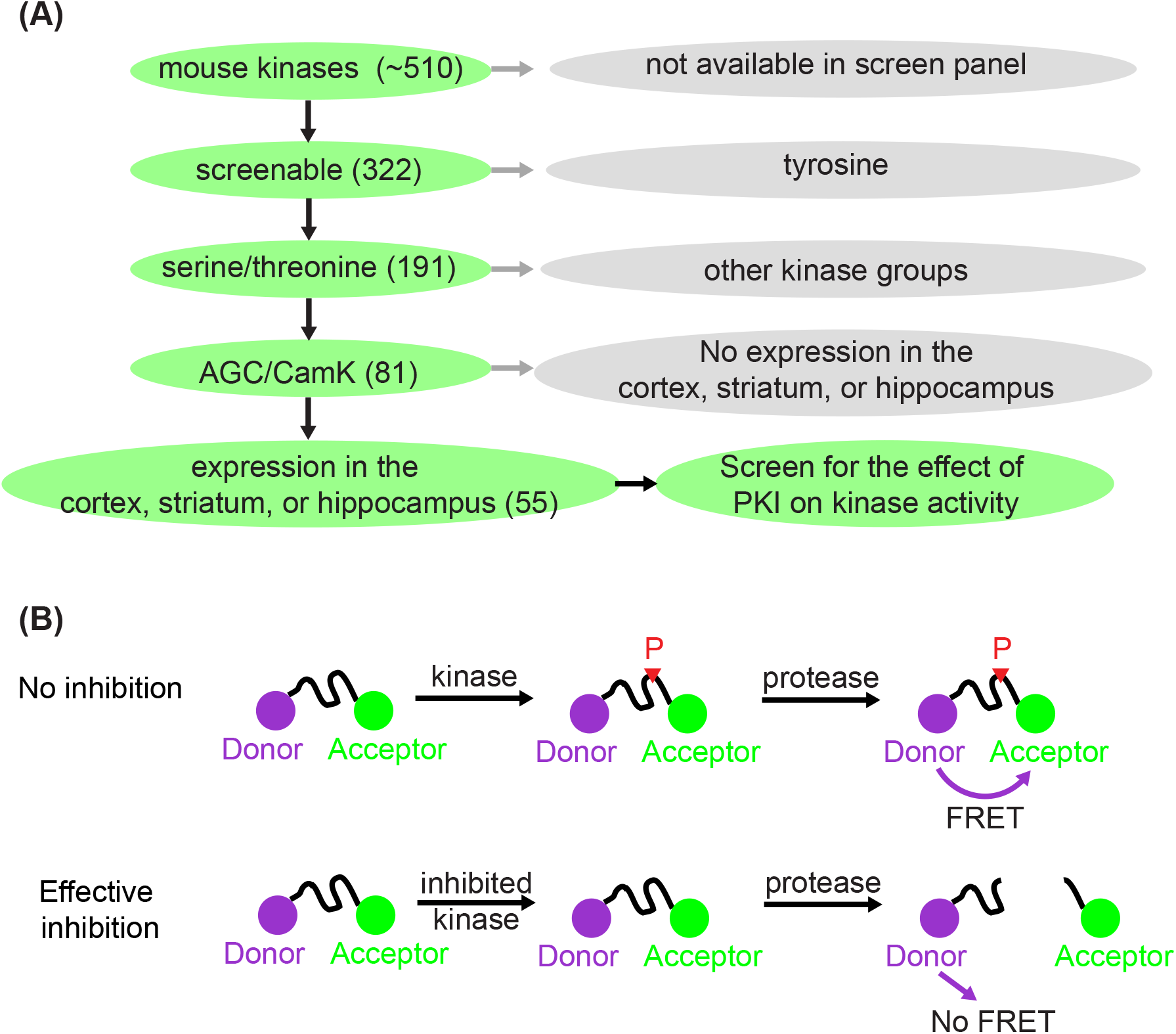
Kinase selection and screening strategy for the effect of PKI on kinases. **(A)** Schematic showing how 55 kinases were selected out of ~510 mouse kinases for screening of the effect of PKI on kinase activity. **(B)** Schematic illustrating the principle of the Z’-LYTE Kinase Assay from Thermo Fisher (Rodems et al., 2002). With no inhibition, the kinase phosphorylates the kinase substrate that separates between the donor (coumarin) and acceptor (fluorescein). The phosphorylation prevents subsequent cleavage by a site-specific protease. Thus, the uncleaved phosphorylated product will exhibit Förster Resonance Energy Transfer (FRET). With effective kinase inhibition, the peptide is not phosphorylated and is subsequently cleaved by the protease, and hence there is no FRET.

### Data analysis

Dose-response curves were fit using variable-slope nonlinear regression (4 parameters) in GraphPad Prism (GraphPad Software). All figures show mean±SEM. Kruskal-Wallis followed by Dunn’s Multiple Comparison Test was used to evaluate statistical significance in Supplementary Figure 1.

## Results

### Kinase screen to determine the specificity of PKI

To investigate whether PKI has effects on kinases beyond PKA, we screened the effects of PKI on a panel of kinases. There are about 510 kinases encoded in the mouse genome, and screening all of them would be too costly. Therefore, we limited our selection of kinases to screen via a tiered system (Figure 1A). First, among the 510 kinases, 322 can be screened by activity enhancement or inhibition via either Adapta Universal Kinase Assay or the Z’-LYTE Peptide Kinase Assay. These are *in vitro*, cell-free assays using purified kinases and therefore suitable for measuring direct kinase action. Second, since PKA is a serine/threonine kinase, we deduced that PKI would most likely target other serine/threonine kinases. Thus, we limited our screen to the 191 serine/threonine kinases among the 322 kinases that can be screened. Third, among the 191 serine/threonine kinases, we reasoned that PKI would mostly act on two kinase groups: the AGC kinases of which PKA is a member, and the CamK kinases since these share substrate similarities with PKA. In total, among the 191 serine/threonine kinases, 81 kinases belong to the AGC and CamK groups. Finally, since PKI plays important roles in neuronal function by regulating synaptic plasticity, and small peptide of PKI has been extensively used as pharmaceutical agent to inhibit PKA activity in the cortex, striatum, and hippocampus, we further narrowed down to 55 kinases that are expressed in these three brain regions based on RNA sequencing data (Cembrowski et al., 2016; Saunders et al., 2018; Zeisel et al., 2018). Therefore, we screened the effects of PKI on 55 kinases, which were selected based on the availability of kinase screen, kinase groups, and expression pattern of these kinases.

Fifty-two kinases were screened with the Z’-LYTE Peptide Kinase Assay (Figure 1B) (Rodems et al., 2002) and three kinases were screened with the Adapta Universal Kinase Assay. In the Z’-LYTE assay, the kinase substrate is flanked by a donor and acceptor fluorophore. Phosphorylation of the substrate prevents subsequent cleavage by a site-specific protease, and thus enables Förster Resonance Energy Transfer (FRET) between the two fluorophores. In contrast, kinase inhibition results in no phosphorylation, and the substrate can therefore be cleaved by the protease, resulting in no FRET. The degree of FRET is therefore correlated with kinase activity. For three of the 55 kinases (CamK1, death-associated protein kinase 1 (DAPK1), and AMPK-related protein kinase 5 (ARK5)), Z’-LYTE assays were not available, and Adapta Universal Kinase Assay was used instead. Adapta is a fluorescence-based immunoassay that detects ADP produced by kinase activity. In summary, two cell-free kinase assays were used to screen for the effect of PKI on fifty-five serine/threonine kinases.

### The effect of PKI (6-22) amide on a panel of kinases

PKI (6-22) amide was used to for the initial screen because this short peptide is a potent inhibitor of PKA and has been widely used to inhibit PKA activity in numerous studies. We first determined with a high concentration of PKI (6-22) amide (5 μM) the kinases whose activities are PKI sensitive at all and therefore merit further studies with dose response curves. As expected, 5 μM PKI (6-22) amide effectively inhibited PKA activity by 85%. PKI (6-22) amide had no effect on the majority of other kinases. However, at 5 μM, PKI (6-22) amide inhibited 5 other kinases by more than 10%. Surprisingly, PKI (6-22) amide (5 μM) enhanced the activity of 20 kinases by more than 10% (expressed in Figure 2 as negative inhibition). Notably, the 8 most facilitated kinases were all isoforms of the PKC family. The amount of facilitation ranged from 59% to 93% (Figure 2).

**Figure 2.**
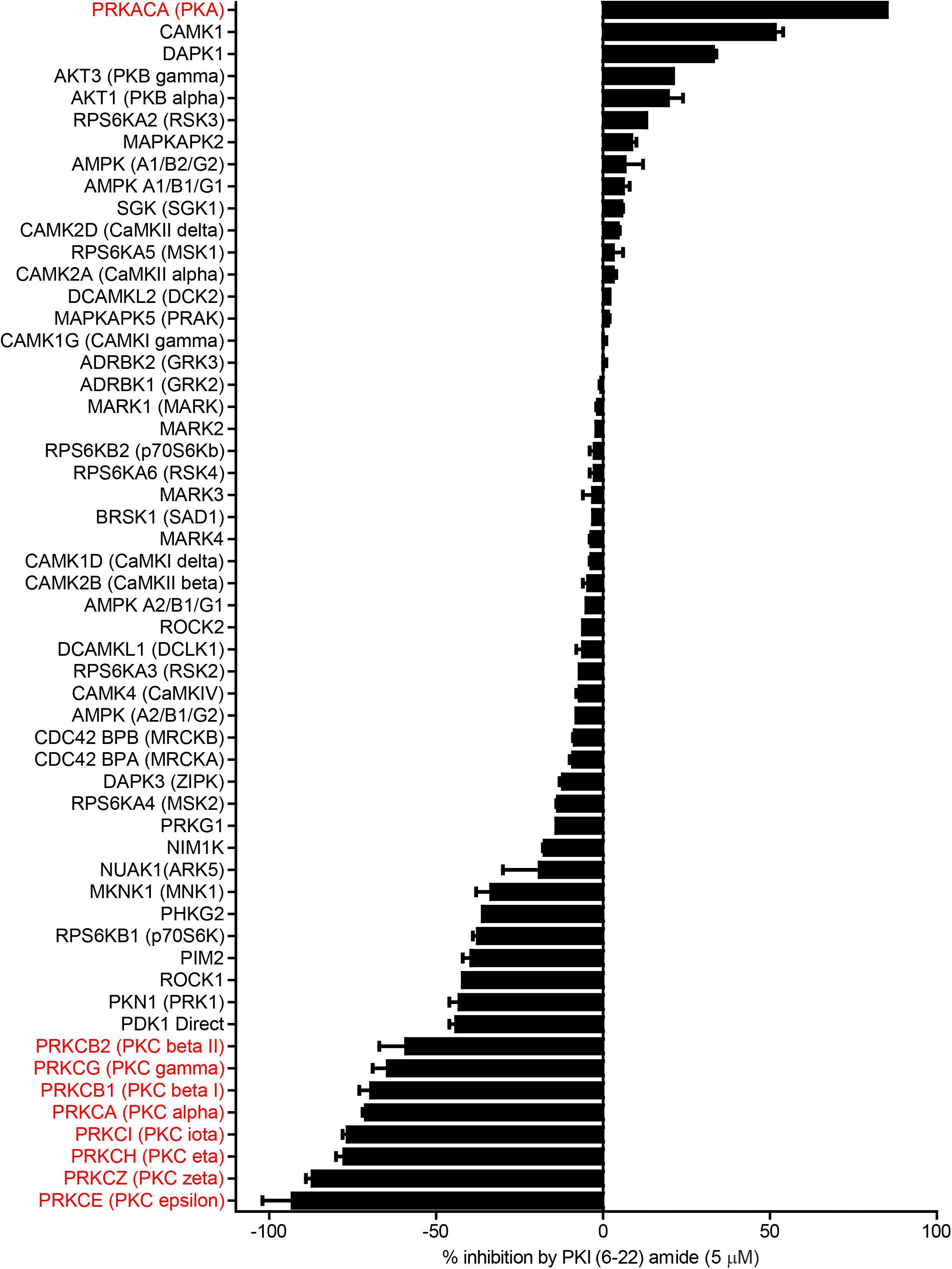
PKI (6-22) amide (5 μM) inhibits some kinases but facilitates others. This bar graph shows the % inhibition of 55 kinases by PKI (6-22) amide (5 μM). The kinases are sorted from the most inhibited (top) to the most facilitated (bottom). The graph shows mean and SEMs.

Since we were surprised by the facilitating effect of PKI (6-22) amide on the PKC family, we set out to disambiguate the potential effect of PKI (6-22) amide on PKC in the kinase reaction, on the protease reaction (development reaction), or on fluorescence itself. PKI (6-22) amide did not induce any significant fluorescence readout change when added in or after the development reaction, but did induce a large change when added in the kinase reaction for PKC epsilon (Supplemental Figure 1). These control experiments indicate that PKI (6-22) amide indeed facilitated PKC activity.

To determine the concentration at which PKI (6-22) amide exerts effects, the dose-response curves of PKI (6-22) amide on activities of 11 selected kinases were measured. These kinases included 1) PKA, 2) all five other kinases whose activity is inhibited by more than 10% at 5 μM, and 3) five selected kinases whose activity is increased by more than 10% at 5 μM – three of these are PKC isoforms. The IC50 of PKI (6-22) amide for PKA was 0.61 nM, comparable to the value determined from previous studies (Glass et al., 1989a) (Figure 3A). PKI (6-22) amide altered the other kinase activities at much higher concentrations (Figure 3A,3B). However, at the PKI concentrations often used in pharmacology experiments (1-10 μM), there were clear effects on several kinases. For example, at 10 μM, PKI (6-22) amide facilitated PKCα and PKCζ by more than 50%. Even at 1.7 μM, PKI (6-22) amide inhibited CamK1 activity by 30%, and facilitated PKCα activity by 33%.

**Figure 3.**
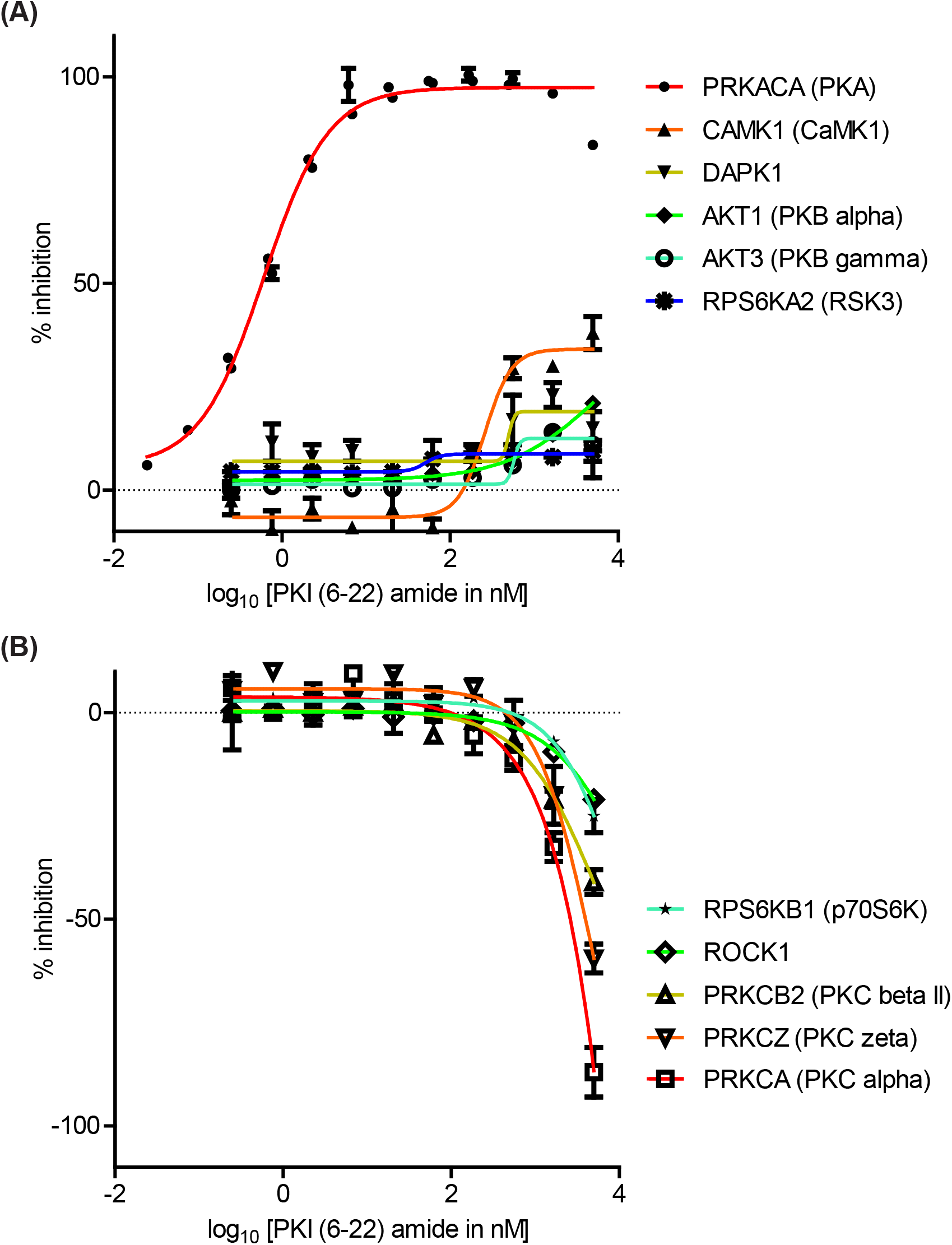
Dose-response curves of the effects of PKI (6-22) amide on kinase activity. **(A)** Dose-response curve of PKI (6-22) amide on inhibited kinases. **(B)** Dose response curve of PKI (6-22) amide on facilitated kinases. The points on the curves show mean and SEMs.

### The effect of full length PKIα on kinase activities

Full length PKI may display different potency and specificity compared with the short peptide. Therefore, although the screen with PKI (6-22) amide above was useful in determining the specificity of PKI as pharmaceutical agents, it is important to perform dose-response curves with full length PKI to assess whether endogenous PKI may alter activities of kinases other than PKA. Full length mouse PKIα protein was synthesized and purified by High-Performance Liquid Chromatography (HPLC), and was subsequently used to determine the dose response curve of 7 kinases (Figure 4).

**Figure 4.**
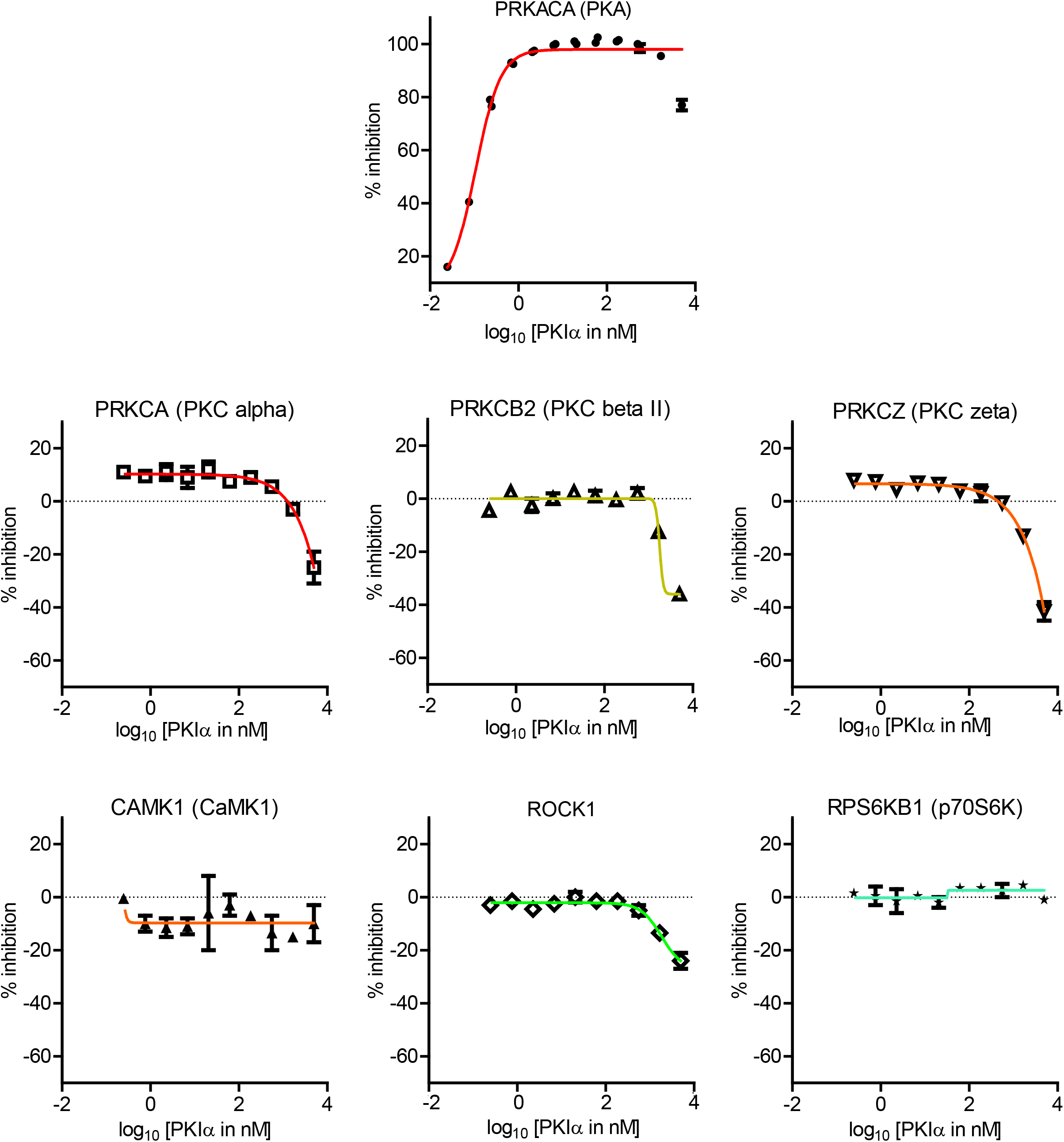
Dose-response curves of the effects of full length PKIα on kinase activity. Dose-response curve of full length PKIα on the activities of 7 kinases. Note a negative value in % inhibition means PKI facilitates kinase activity. The points on the curves show mean and SEMs.

PKIα inhibited PKA activity with an IC50 of 0.11 nM, with higher potency than PKI (6-22) amide (Figure 4). PKIα did not have significant effect on CamK1 or p70S6K (Figure 4). In addition, PKIα also facilitated multiple PKC isoforms and ROCK1 at much higher concentrations (1-5 μM) (Figure 4). At 5 μM, full length PKIα facilitated PKCα, PKCβII, PKCζ, and ROCK1 by 25%, 36%, 42%, and 24% respectively.

## Discussion

In order to understand the specificity of PKI on kinases, we screened both PKI (6-22) amide and full length PKIα against a panel of 55 kinases. In addition to inhibiting PKA activity, PKI (6-22) amide also inhibited CaMK1, and facilitated the activities of p70S6K, ROCK1, and multiple PKC isoforms. Full length PKIα, at high concentrations, also facilitated the activity of ROCK1 and multiple PKC isoforms. These results inform the design and interpretation of experiments using PKI as a pharmacological agent, provide a starting point to explore novel functions of endogenous PKI, and raise the interesting possibility that PKI may act as a molecular switch between PKA and PKC activity.

### Facilitating effects of PKI on PKC

We found that in addition to inhibiting PKA activity, PKI also facilitated the kinase activity of multiple PKC isoforms. This was surprising because previous studies found little influence of PKI on PKC (Day et al., 1989; Smith et al., 1990). In principle, this could be due to PKI interfering with 1) the protease reaction, 2) the fluorescence detection, or 3) the FRET response. Interference with the protease reaction was minimal, because PKI had very little effect on the control assay with unphosphorylated cleavable substrates (PKI+no ATP versus no ATP conditions in the kinase reaction). In addition, adding PKI in the protease development reaction for PKC did not result in a significant FRET change (Supplementary Figure 1). Similarly, neither full length PKIα nor PKI (6-22) amide is autofluorescent, as evidenced by the test compound fluorescence interference control with no kinase/peptide mixture. Finally, PKI did not have any significant effects on FRET itself, because the FRET readout was not significantly different when PKI was added after the development reaction. These data indicate that PKI facilitated the kinase activity of multiple PKC isoforms.

Several technical differences may explain the differences between our results and previous studies that did not find significant influence of PKI on PKC. First, these studies differed in the form of PKI used: Smith et al. (Smith et al., 1990) used PKI-tide whose sequence is **IA<u>A</u>GRTGRR<u>Q</u>AI**HDILVAA, whereas we used PKI (6-22) amide whose sequence is TYADF**IA<u>S</u>GRTGRR<u>N</u>AI**(bolded amino acids are the shared fragment, and differences in sequences within the shared fragment are underlined). Second, the forms of PKC used are different: we used purified human recombinant PKC isoforms expressed in insect cells, whereas previous studies used partially purified PKC without clear distinction of isoforms (Day et al., 1989; Smith et al., 1990). Finally, although previous results did not reach statistical significance, Day et al. (Day et al., 1989) showed a mild enhancement of PKC activity by full length PKI in a cell-based assay with PKI transfection.

### Interpreting pharmacology experiments with PKI

PKI (6-22) amide and other short peptides of PKI are widely used in pharmacological experiments to inhibit PKA activity. They are often used at between 1 and 10 μM, either intracellularly through a whole-cell recording pipette, or applied in the bath in cell-permeant myristoylated forms. Here, we found that PKI (6-22) amide, at these concentrations, also inhibits CamK1, and facilitates the activity of multiple PKC isoforms, possibly acting as a positive allosteric modulator. Thus, these additional targets of PKI need to be taken into account to re-interpret previous results using PKI, and to design and interpret future experiments with PKI.

### Significance of PKI effects for biology

We found that full length PKIα, one of the endogenously expressed isoforms, enhanced the activity of ROCK1 and multiple PKC isoforms at high concentrations. This raises the interesting possibility that PKI acts as a molecular switch to regulate the balance between PKA and PKC activity. However, the validity of this hypothesis depends on the endogenous concentration of PKIα, which is hard to determine, partially because the local concentrations of PKIα within a subcellular compartment may be higher than what can be determined with traditional biochemical methods. Our study therefore lays the foundation for future explorations on how endogenous PKI impacts cellular signaling and confines it in space and time through non-classical targets.

## Author contributions

Y.C. designed the experiments, implemented the experiments, analyzed the data, and wrote the manuscript. B.L.S. provided feedback on experimental design and data analysis, and edited the manuscript.

## Acknowledgments

We thank the team at ThermoFisher SelectScreen Kinase Profiling Services for helpful discussions and additional control experiments. This work was supported by grants from the Brain & Behavior Research Foundation (NARSAD Young Investigator Grant 28323 to Y.C.), a Goldenson Fund postdoctoral fellowship (to Y.C) and NINDS (R37NS046579, to B.L.S.)

