## Supplementary Figure 1 for "The kinase specificity of protein kinase inhibitor peptide (PKI)"

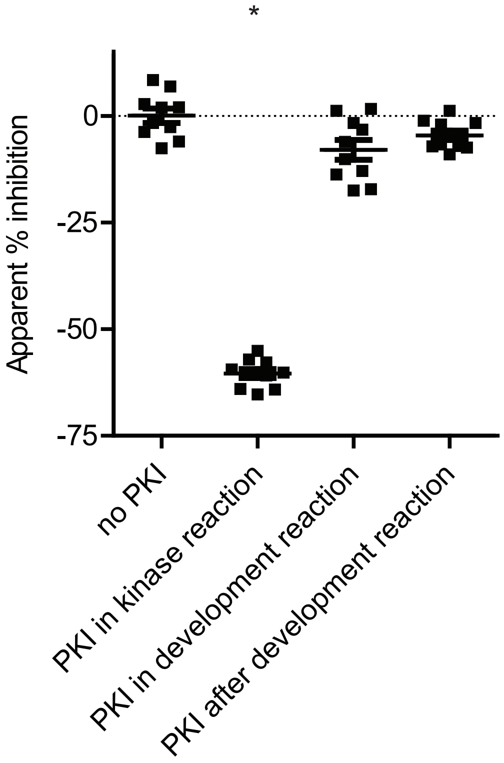


**Supplementary Figure 1. PKI (6-22) amide inhibits kinase reaction without significant effect on the protease/development reaction or fluorescence of the substrate.** Apparent % inhibition, based on FRET ratio, of adding PKI in the kinase reaction of PRKCE (PKC epsilon), the protease development reaction, or after development reaction. *: $p<0.05$ vs. no PKI, $p<0.05$ vs. PKI in development reaction, $p<0.05$ vs. after development reaction (Kruskal-Wallis followed by Dunn's Multiple Comparison Test). The graph shows mean and SEMs.
